# Genetic variation and RNA structure regulate microRNA biogenesis

**DOI:** 10.1101/093179

**Authors:** Noemi Fernandez, Ross A. Cordiner, Robert S. Young, Nele Hug, Sara Macias, Javier F. Cáceres

## Abstract

MiRNA biogenesis is highly regulated at the post-transcriptional level; however, the role of sequence and secondary RNA structure in this process has not been extensively studied. A single G to A substitution present in the terminal loop of pri-mir-30c-1 in breast cancer patients had been previously described to result in increased levels of mature miRNA. Here, we report that this genetic variant directly affects Drosha-mediated processing of pri-mir-30c-1 in *vitro* and in cultured cells. Structural analysis of this variant revealed an altered RNA structure that facilitates the interaction with SRSF3, an SR protein family member that promotes pri-miRNA processing. Our results are compatible with a model whereby a genetic variant in pri-mir-30c-1 leads to a secondary RNA structure rearrangement that facilitates binding of SRSF3 resulting in increased levels of miR-30c. These data highlights that primary sequence determinants and RNA structure are key regulators of miRNA biogenesis.

---

MicroRNAs (miRNAs) are short non-coding RNAs that negatively regulate the expression of a large proportion of cellular mRNAs, thus affecting a multitude of cellular and developmental pathways^1,2^. The canonical miRNA biogenesis pathway involves two sequential processing events catalyzed by RNase III enzymes. In the nucleus, the Microprocessor complex, comprising the RNase III enzyme Drosha, the double-stranded RNA-binding protein, DGCR8, and additional proteins carries out the first processing event, which results in the production of precursor miRNAs (pre-miRNAs)^3,4^. These are exported to the cytoplasm, where a second processing event carried out by another RNase III enzyme, Dicer, leads to the production of mature miRNAs that are loaded into the RISC complex^5^.

Due to the central role of miRNAs in the control of gene expression, their levels must be tightly controlled. As such, dysregulation of miRNA expression has been shown to result in grossly aberrant gene expression and leads to human disease^6–9^. In particular, the Microprocessor-mediated step of miRNA biogenesis has been shown to be regulated by multiple signaling pathways, including the transforming growth factor-β (TGF-β) pathway as well as the p53 response, leading to activation of subsets of individual miRNAs^10,11^. Furthermore, altered miRNA expression has been associated with the progression of cancer^12,13^, where a global downregulation of miRNA expression is usually observed^14,15^. It was recently shown that miRNA biogenesis can also be regulated in a cell-density-dependent manner via the Hippo-signaling pathway, and that the observed perturbation of this pathway in tumors may underlie the widespread downregulation of miRNAs in cancer^16^. Thus, miRNA production is tightly controlled at different levels during the biogenesis cascade. Extensive evidence has shown that RNA-binding proteins (RNA-BPs) recognize the terminal loop of miRNA precursors and influence either positively or negatively the processing steps carried out by Drosha in the nucleus and/or Dicer in the cytoplasm. These include the hnRNP proteins, hnRNP K and hnRNP A1, as well a the cold-shock domain protein, Lin28 and the RNA helicases, p68/p72^17^. In the case of the multifunctional RNA-binding protein hnRNP A1, we have previously shown that it can act as an auxiliary factor by binding to the conserved terminal loop of pri-miR-18a and promote its Microprocessor-mediated processing^18,19^. Conversely, the same protein can act as a repressor of let-7 production in differentiated cells^20^.

Several studies have shown that there is a correlation between the presence of polymorphisms in pri-miRNAs and the corresponding levels of mature miRNAs^21^; however, a mechanistic understanding of how sequence variation and RNA structure control miRNA biogenesis has not been explored in great detail. Screening of novel genetic variants in human precursor miRNAs linked to breast cancer identified two novel rare variants in the precursors of miR-30c and miR-17, resulting in conformational changes in the predicted secondary structures and also leading to altered expression of the corresponding mature miRNAs. These patients were non-carriers of BRCA1 or BRCA2 mutations, suggesting the possibility that familial breast cancer may be caused by variation in these miRNAs^22^. In particular, the single G to A substitution in primary miR-30c-1 (pri-mir-30c-1) terminal loop, which was also later observed in gastric cancer patients^23^, results in an increase in the abundance of the mature miRNA.

Here, we investigated the mechanism by which the pri-mir-30c-1 variant detected in breast cancer patients results in an increased expression of this miRNA. We found that this genetic variant directly affects the Microprocessor-mediated processing of this miRNA. A combination of structural analysis with RNA chromatography coupled to Mass spectrometry revealed changes in the pri-miRNA structure that lead to differential binding of a protein factor, SRSF3, that has been previously reported to act as a miRNA biogenesis factor. These results provide a mechanism by which the pri-mir-30c-1 genetic variant results in an increased expression of the mature miR-30c. Altogether these data highlights that primary sequence as well as RNA structure have a crucial role in the post-transcriptional regulation of miRNA biogenesis.

## RESULTS

### The G/A substitution in pri-miR-30c-1 affects the Microprocessor-mediated processing of the primary miRNA

In order to understand the mechanism underlying miR-30c deregulation in breast and gastric cancers, we investigated how the reported G_27_-to-A mutation observed in a Chinese population might affect miRNA biogenesis. It was previously shown that this substitution results in an increase in the abundance of the mature miRNA; however, the mechanism that leads to an increased expression is unknown^22,23^. First, we transiently transfected MCF7 breast cancer cells with constructs encoding 380 nucleotides (nt) of primary hsa-pri-mir-30c-1 (pri-miRNA), either in a WT version or bearing the G/A variant (**Fig. 1a**). We observed that the G/A substitution resulted in increased levels of mature miR-30c (**Fig. 1b**), resembling the situation observed in patients with this mutation. Furthermore, this was not due to increased transcription of the G/A-harboring pri-miRNA, as shown by unchanged levels pri-miRNA levels (data not shown). In order to dissect the precise step of miRNA biogenesis pathway that is affected by the G/A substitution, we used an RNA version of pri-mir-30c-1 that has yet to undergo processing by the Microprocessor in the nucleus and by Dicer in the cytoplasm (pri-miRNA). As a counterpart, we transfected an RNA oligonucleotide that mimics the precursor miRNA (pre-miRNA), a sequence that arises upon processing by the Microprocessor. Importantly, we observed approximately a two-three fold increase of miR-30c mature levels when transfecting the G/A sequence derived from the pri-miRNA sequence, whereas no changes were detected following transfection of the pre-miRNA sequence (**Fig. 1c**). This experiment demonstrates that the G/A substitution exclusively affects the Drosha-mediated processing of the pri-miRNA. Moreover, we could recapitulate this result in an *in vitro* reaction. We found that *in vitro* transcribed pri-mir-30c-1 was readily processed in the presence of MCF7 total extracts, rendering a product of ~65 nts that corresponds to pre-mir-30c. Notably, the processing of the G/A variant was increased, when compared to the WT version, as was observed in living cells (**Fig. 1d**). The effect of the G/A variation in the processing of pri-mir-30c-1 was also recapitulated using a purified Microprocessor (FLAG-Drosha/FLAG-DGCR8 complex) and a shorter *in vitro* transcribed substrate (153nt) **(Supplementary Fig. 1)**. Altogether, these complementary approaches indicate an enhanced Microprocessor-mediated processing of the G/A variant sequence and this recapitulates what was previously observed in breast and gastric cancer patients.

**Figure 1.**
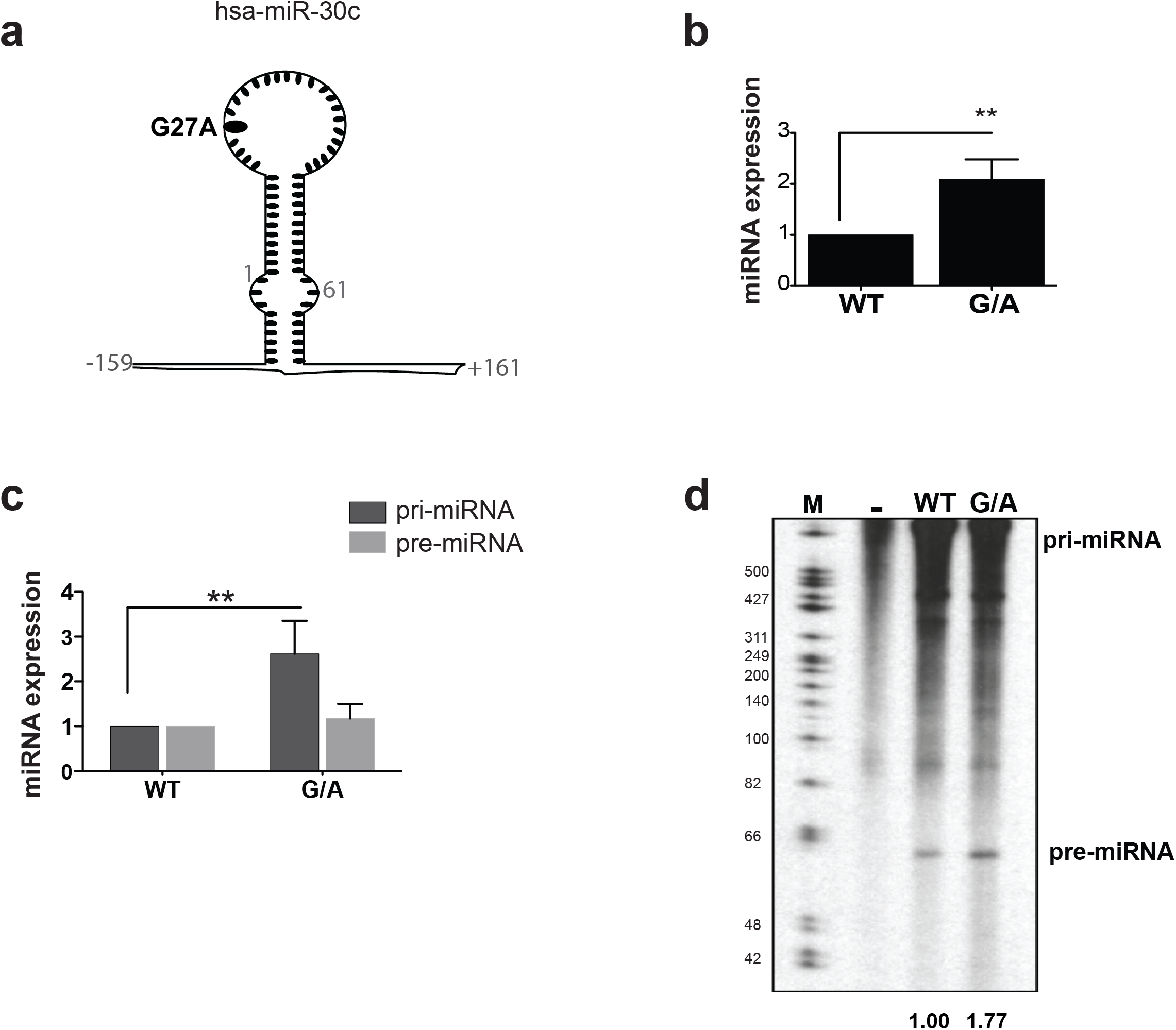
A genetic variant in the terminal loop of hsa-pri-mir-30c-1 alters its normal expression. **(a)** Schematic representation of the hsa-pri-mir-30c-1 transcript indicating the G to A mutation observed in breast and gastric cancer patients. Nucleotides (nt) encompassing primary (pri-miRNA; nt −159 to +161) and precursor miRNA (pre-miRNA; nt 1 to 61) used in this experiment are indicated. Numbers are relative to the first nt of the mature miRNA. **(b)** The relative expression levels of mature miR-30c in MCF7 cells transfected with plasmids encoding either a wild-type sequence (WT) or one bearing the G/A mutation (n=12) **(c)** Levels of mature miR-30c in MCF7 cells transfected with *in vitro* transcribed pri-mir-30c-1 (pri-miRNA) or an RNA oligonucleotide that mimics pre-miRr-30c-1 (pre-miRNA) sequences, either in a wild-type or G/A version (n=5). Mann-Whitney U test was used to evaluate differences between WT and G/A samples. Error bars indicate SEM. ** p<0.01. **(d)** Representative *in vitro* processing of pri-mir-30c-1 (380 nt) in MCF7 total cell extracts. Quantification of pre-miRNA band intensities are shown below and expressed as the relative intensity normalized to pre-miR-30c-1 WT variant.

### The sequence of the terminal loop in pri-miR-30c-1 is highly conserved across species

Experiments described above confirmed a crucial role for the G_27_ residue in influencing miR-30c biogenesis. We analyzed the genomic variation in the hsa-pri-mir-30c-1 sequence across vertebrates, and detected substantial evolutionary constraint across the entire locus, as indicated by positive GERP (Genomic Evolutionary Rate Profiling) scores (**Fig. 2a**). Constrained residues, which highlight regions under purifying selection, are located in the mature miRNA sequences in both arms, as expected due to their effect in the regulation of gene expression. Interestingly, part of the terminal loop (TL), where G_27_ is embedded, has also a very high level of constraint, which is suggestive of a role of this sequence in miRNA biogenesis, as was previously described for a subset of miRNAs^17,19,24^. In addition, residues at the 5´ end (nts −1 to −8; −11 to −16 and −20 to −23) and the 3´end (nts +1 to +16 and +18 to +25) are also highly constrained. Indeed, several of these residues were included as part of the stem in the *in silico* predicted RNA structure suggesting their importance for maintaining the RNA secondary structure (**Fig. 2b**). These data led us to focus our attention on these invariant sequences as potentially having a crucial role in the regulation of miR-30c biogenesis.

**Figure 2.**
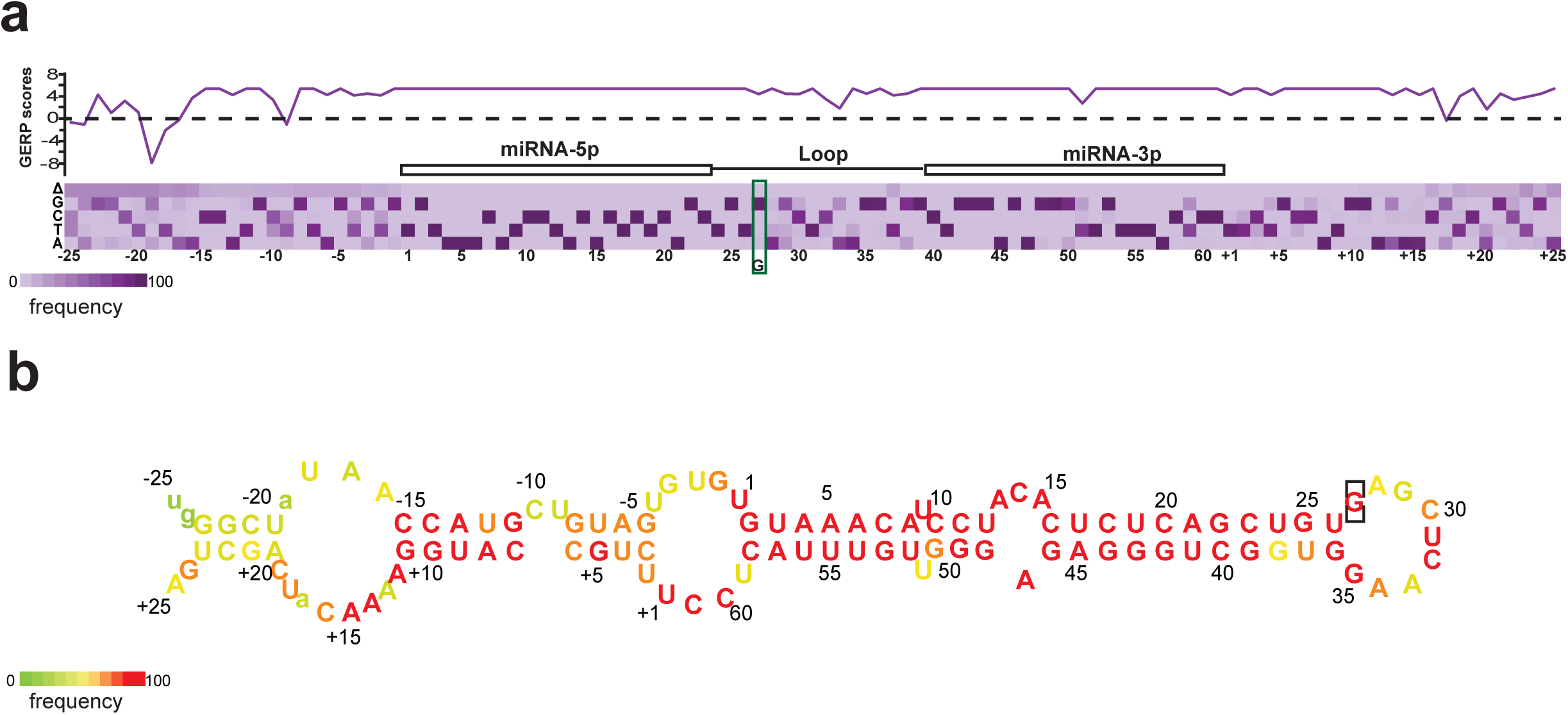
Sequence conservation of pri-mir-30c-1 precursors **(a)** Nucleotide-level GERP scores across the locus, indicating extensive evolutionary constraint. A GERP score above zero indicates significant constraint, while a score below zero indicates an excess of nucleotide substitutions beyond the expected neutral rate. The purple bars display the total number of observed nucleotide substitutions found in aligned sequences from 98 vertebrates. Δ represents absence of the nucleotide from 98 vertebrate sequences analyzed. The location of the modified G/A nucleotide is indicated by a green rectangle. **(b)** Model of the predicted secondary structure with nucleotides colored as in (a) to reflect their variation. Nucleotides present in less than 50% of the species are indicated in lower case.

### RNA structural analysis reveals the specific requirement of G_27_ for maintaining the pri-miRNA structure

Next, in order to establish the importance of the G_27_ to A substitution in RNA structure, we performed structural analysis by Selective 2’-hydroxyl acylation analyzed by primer extension (SHAPE)^25^. This approach allows performing quantitative RNA structural analysis at single nucleotide resolution and is mostly independent of base composition. While highly reactive residues are located at single-stranded regions, non-reactive nucleotides are involved in base pairs, non-Watson-Crick base pairs, tertiary interactions or single stacking interactions in the C2´-endo conformation^26^. To this end, *in vitro* transcribed RNA comprising 380 nt of pri-miR-30c-1 (either wild-type or the G/A variant sequence) was treated with N-methylisatoic anhydride (NMIA), which reacts with the 2´hydroxyl group of flexible nucleotides (**Fig. 3a**). Gross modifications of SHAPE reactivity were observed in specific regions of the G/A variant, when compared with the wild-type sequence. The resulting profiles revealed a decreased SHAPE reactivity in the TL (residues 28-30), with a concomitant increase in the 5' region (−18,−16, and −15) as well as in the 3´end (nts +11, +16 and +19) (**Fig. 3a,b**). This result indicated the presence of different conformations in the pri-miRNA with the G-to-A substitution, as compared to its wild-type counterpart. In order to gain more information into the folding and tertiary structure of this pri-miRNA, we assessed the solvent accessibility of each nucleotide by hydroxyl radical cleavage footprinting, generated by reduction of hydrogen peroxide by iron (II)^27^. Hydroxyl radicals break the accessible backbone of RNA with no sequence dependence. We defined buried regions, as zones with more than two consecutives nucleotides having a reactivity (R) smaller than the mean of all reactivity, whereas exposed regions are those with more than two consecutive nucleotides having R larger than the mean of all reactivity. We observed that the wild-type sequence presents two buried regions located between nts 8 to 40 and +9 to +25, as well as two exposed segments between nt −25 to 7 and nts 40 to +8 **(Supplementary Fig. 2a)**. The G to A substitution caused changes in the exposure to solvent, with both the TL (nts 28 to 40) and also the 3´end region (nts +17 to +22) becoming solvent accessible. By contrast, the 5´end (nts −7 to 7) and a small region in miRNA-3p (nts 55 to 58) are no longer solvent accessible.

**Figure 3.**
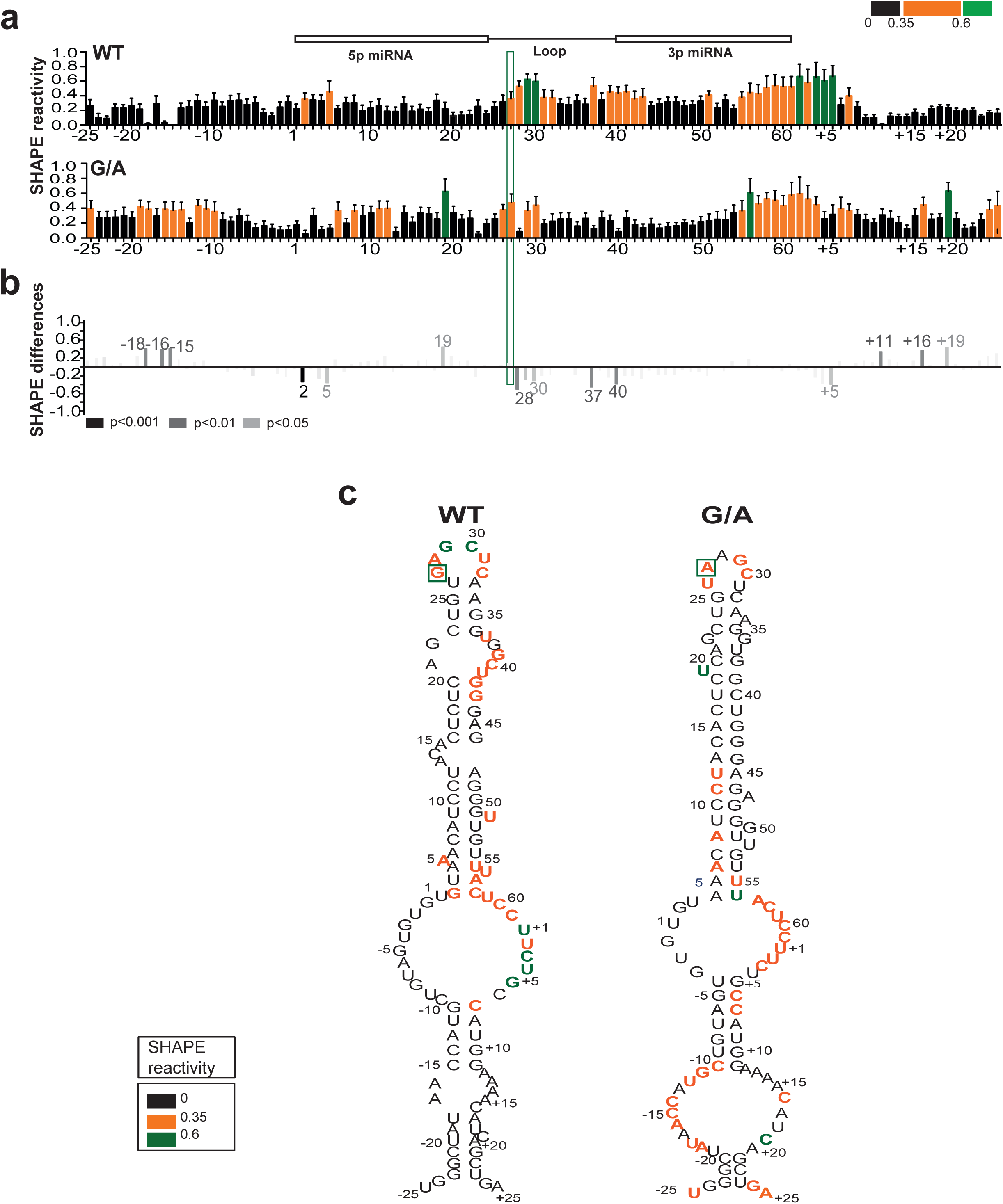
Structural analysis reveals the importance of the G to A substitution in pri-miR-30c-1 **(a)** Values of SHAPE reactivity (depicted using color coded bars) at each individual nucleotide position correspond to the mean SHAPE reactivity (+/− SEM) (n=10). **(b)** SHAPE differences plots of the G/A variant relative to the wild-type sequence. Only nucleotides with absolutes differences bigger than 0.25 and statically significant are depicted (grey colors represent p-values). Bars above the basal line indicate increased SHAPE reactivity for the G/A variant, whereas those below the basal line indicate lower SHAPE reactivity. Mann-Whitney U test was used to evaluate differences between WT and G/A samples. **(c)** Predicted secondary structures of WT and G/A variant sequences, as determined by SHAPE reactivity values.

Altogether, the SHAPE and radical hydroxyl data suggest that the G_27_A substitution is indeed affecting the RNA flexibility of pri-miRNA-30c-1, modifying both base-pairing interaction as well as solvent accessibility of the nucleotides located in the TL and in the basal region of the stem. This could be a consequence of a long distance interaction disruption between those regions (**Fig. 3c** and **Supplementary Fig. 2b**), which could in turn modify the interaction with RNA-binding proteins important for miR-30c biogenesis.

### SRSF3 binds to a basal region of hsa-pri-mir-30c-1

A working model that emerges from data described above is that either a repressor of Microprocessor-mediated processing binds to the wild-type sequence or, alternatively, the change in RNA structure induced by the G/A sequence variation could lead to the binding of an activator. In order to identify RNA-BPs that differentially bind to either the pri-miR-30c-1 WT or G/A sequence, we performed RNase-assisted chromatography followed by mass spectrometry in MCF7 total cell extracts^28^. This resulted in the identification of twelve proteins that interact with the WT sequence and eight that bind to the G/A variant, being 7 common between both substrates. Significantly, several of the common proteins were previously implicated in miRNA biogenesis and/or regulation, including the heat shock cognate 70 protein^5^, the hnRNP proteins, hnRNP A1^18,20^ and hnRNP A2/B1^29^, as well as the RNA helicase DDX17^4^, and Poly [ADP-ribose] polymerase 1 (PARP)^30^ (**Fig. 4a** and **Supplementary Fig. 3a**).

**Figure 4.**
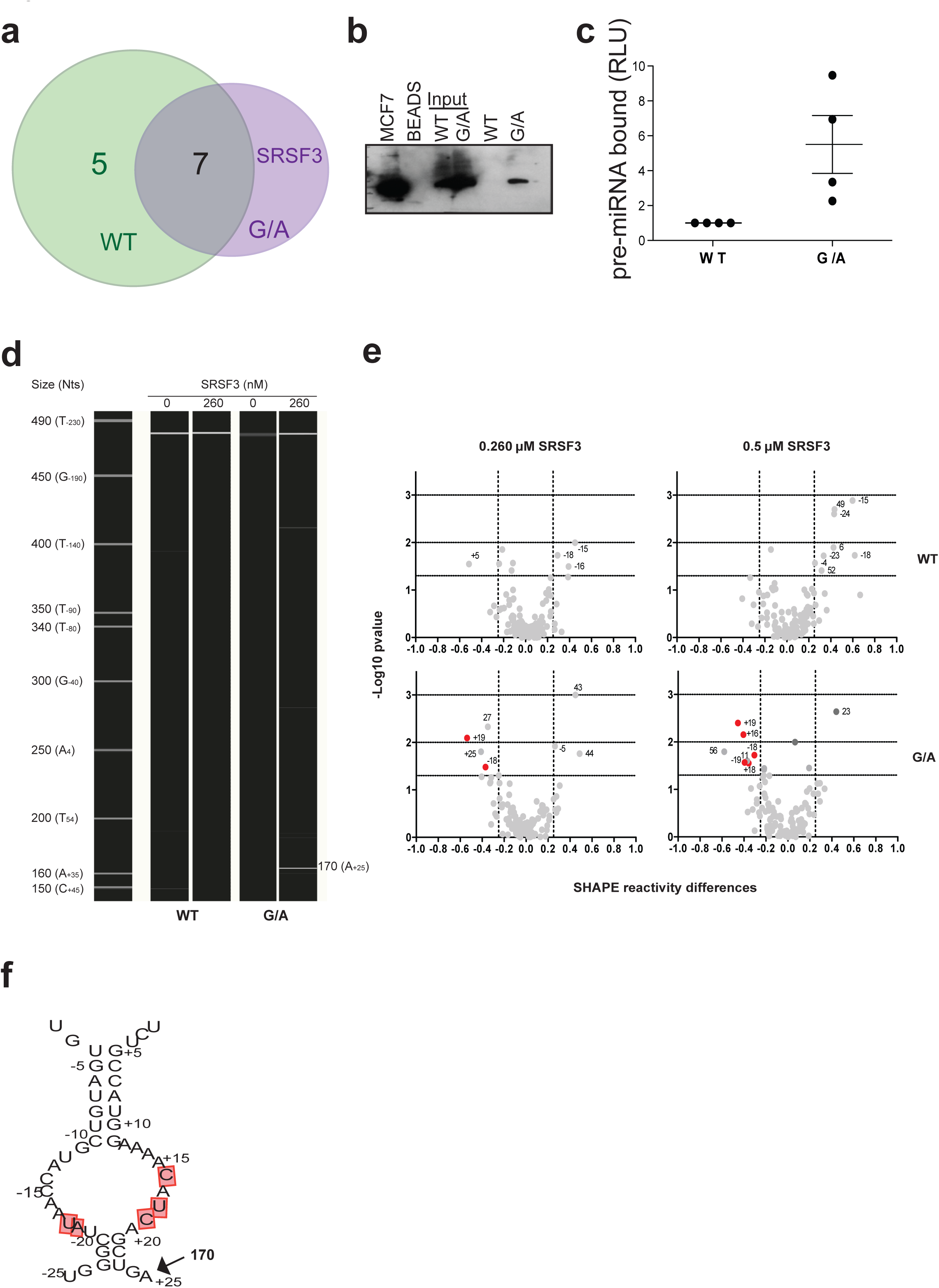
Identification of trans-acting factors binding to pri-mir-30c-1. **(a)** Venn-diagram indicating the number of interactors observed for the WT sequence and the G/A variant. **(b)** Validation of the interaction of SRSF3 with the G/A variant sequence by RNA chromatography followed by Western-blot analysis with an antibody specific for SRSF3. **(c)** Immunoprecipitation with a specific anti-SRSF3 antibody followed by RT-qPCR quantification (+/− SEM) (n=4). **(d)** Representative toeprinting assay in the presence or absence of 260nM of purified SRSF3 protein using fluorescent-labeled primers and capillary electrophoresis. The unique RT stop in the G/A variant (n=2) is indicated. **(e)** Volcano plots showing the SHAPE reactivity data in the presence of 260nM (left panel) or 500 nM (right panel) of purified SRSF3 protein. The red dots indicate those positions protected by SRSF3 in a dose-dependent manner (n=5). Mann-Whitney U test was used to evaluate differences **(f)** Schematic representation of the basal stem of hsa-miR-30c G/A illustrating the nucleotides protected by SRSF3 (in red) as well as the RT stop detected by toeprinting assay.

We decided to focus on those interactors that showed preferential interaction with either the WT or the G/A variant sequence. In this category, the most likely candidate that interacted with the WT sequence is the RNA-binding protein FUS/TLS (fused in sarcoma/translocated in liposarcoma). Interestingly, this protein has been shown to promote miRNA biogenesis by facilitating the co-transcriptional recruitment of Drosha^31^; however, we could not validate the specific interaction between FUS and miR-30c-1 WT sequence by Immunoprecipitation followed by Western blot analysis (IP-WB) **(Supplementary Fig. 3b)**. As for the G/A variant, there was a single exclusive interactor, SRSF3, which is a member of the SR family of splicing regulators. These family of proteins are involved in constitutive and alternative splicing, but some of them have been shown to fulfil other cellular functions^32,33^.

Importantly, SRSF3 has also been reported to be required for miRNA biogenesis^34^. We could validate this interaction by RNA chromatography followed by Western-blot analysis with an antibody specific for SRSF3 (**Fig. 4b**). We also observed preferential binding of endogenous SRSF3 protein to the G/A variant of pri-mir-30c-1, as shown by immunoprecipitation of SRSF3 followed by RT-qPCR quantification of the associated pri-miRNA (**Fig. 4c**). In order to analyze the interaction of SRSF3 with pri-miR-30c-1 wild-type and G/A sequences, we carried out toeprint and SHAPE assays using SRSF3 protein purified from MCF7 cells. Toeprint analysis was performed with fluorescent labeled antisense primers and capillary electrophoresis^35^. In this assay, bound SRSF3 will block the reverse transcriptase and will illuminate the site where SRSF3 is bound to RNA. Prominent toeprint of SRSF3 with the G/A sequence was observed around nt A_+25_-G_+24_ (RNA size 170nt, position A_+25_)(**Fig. 4d**). Similarly, analysis of SHAPE reactivity in the presence of added purified SRSF3, revealed a dose-dependent protection from NMIA attack upon addition of SRSF3 in specific RNA residues in a dose-dependent manner (nts −19, −18, +16,+18,+19) of the basal region of G/A (**Fig. 4f,e**). Of importance, a conserved CNNC motif (nts from +16 to +22), previously described as SRSF3 binding site^34,36,37^ is located within the recognition place. Together, we can conclude that the interaction of SRSF3 with pri-miR-30c-1 takes place at the CNNC motif at the basal region of the G/A variant.

### SRSF3 protein is responsible for the increased levels of the G/A variant of miR-30c

As previously described, SRSF3 was proposed to have a role in miRNA biogenesis by recognition of a CNNC motif located 17nts away from Drosha cleavage site^34^. Pri-mir-30c-1 has two overlapping CNNC motifs (residues from +16 to +21). Notably, accessibility around this region increased in the G/A variant, as determined by SHAPE and hydroxyl radical analysis (**Fig. 3** and **Supplementary Fig. 2**). Furthermore, toeprint and SHAPE assays in the presence of purified SRSF3 protein confirmed the specific recognition of the CNNC motif by SRSF3 in a dose-dependent manner (**Fig. 4**). Next, we addressed whether the preferential binding of SRSF3 to the pri-mir-30c G/A variant sequence was responsible for its increased expression by comparing the mature levels of miR-30c-1 wild-type or G/A variant under variable levels of SRSF3 expression. To this end, we co-transfected pri-mir-30c-1 constructs in MCF7 cells under transiently overexpression of SRSF3, or alternatively, transfected specific siRNAs to knock-down endogenous SRSF3 protein **(Supplementary Fig. 4)**. Of interest, we observed that reduced levels of SRSF3 drastically decreased the levels of the G/A miR-30c variant, without affecting the levels of wild-type miR-30c (**Fig. 5a**, compare WT vs G/A panels). By contrast, transient overexpression of SRSF3 increased significantly the levels of wild-type mature miR-30c, but has a more modest effect on the G/A variant sequence. Altogether, these experiments suggest that SRSF3 binding is limiting in the WT scenario and that is essential to promote miRNA biogenesis in the G/A context.

**Figure 5.**
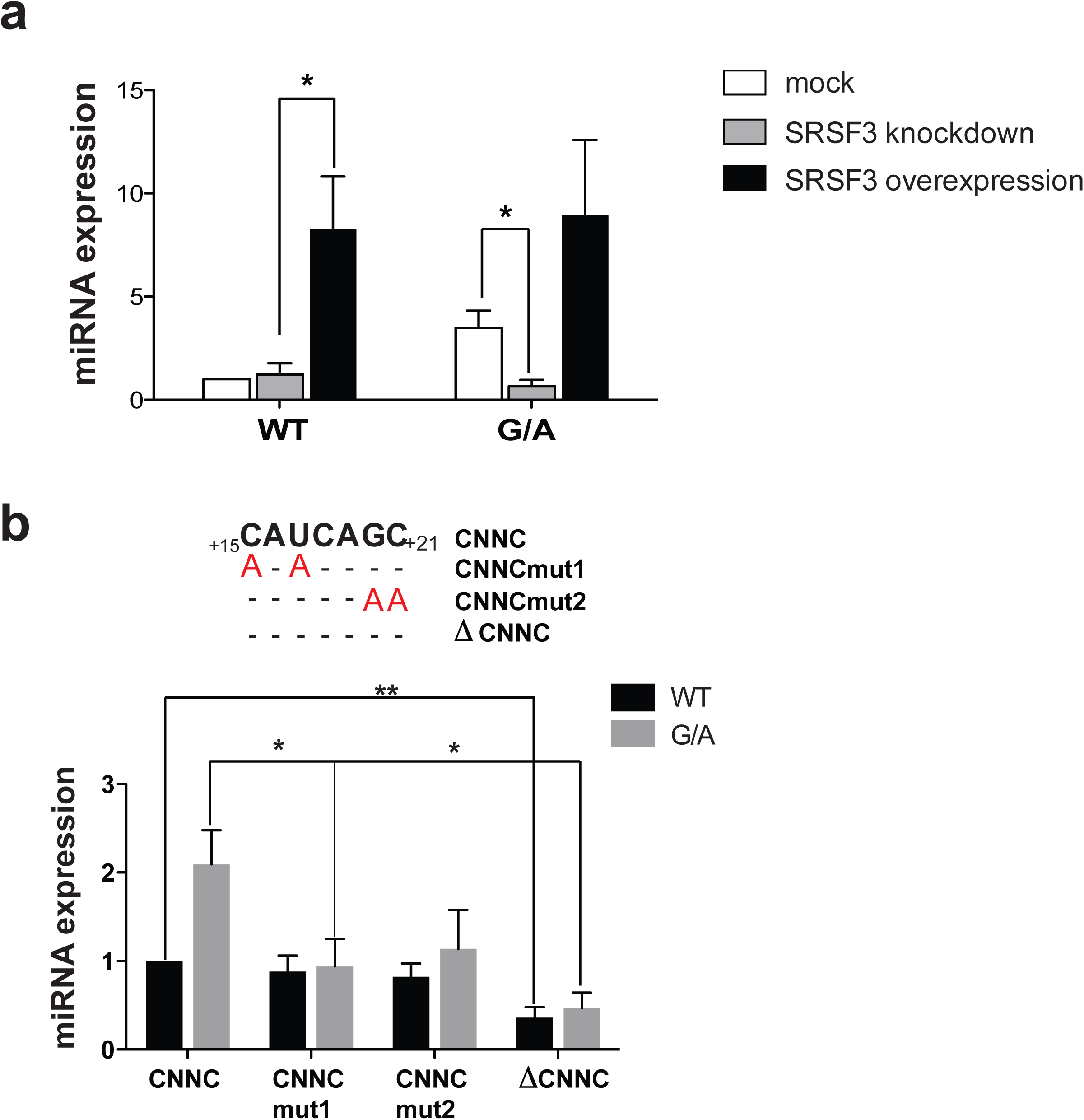
Role of SRSF3 in miR-30c expression. **(a)** The relative expression levels of miR-30c WT and G/A in MCF7 cells with changing expression of SRSF3 protein. Error bars indicate SEM (n=5) * p<0.05 ** p<0.01 **(b)** Mutational analysis of the CNNC motif. Nucleotides substitutions present in the CNNC region are shown in Red. Relative expression levels of miR-30c in MCF7 cells transfected with plasmids encoding either a wild-type sequence (WT) or one bearing the G/A mutation, and also including mutations in the CNNC motif, when indicated. Mann-Whitney U test was used to evaluate differences. Error bars indicate SEM (n=6) * p<0.05 ** p<0.01.

To confirm the role of SRSF3 in the differential processing observed with pri-mir-30c-1 G/A variant sequence, we proceeded to mutate the two consecutive CNNC motifs that are the natural binding sites for SRSF3 (**Fig. 5b**). We generated a set of mutants that affected either the first or second CNNC motif (mut1 and mut2, respectively) or a deletion of both motifs (ΔCNNC). The CNNCmut1 carrying a double substitution C_+15_U_+17_ to AA, led to a severe reduction in the levels of miR-30c expression only with the G/A variant sequence (**Fig. 5b**). Similarly, a double substitution of the second CNNC motif G_+20_C_+21_ to AA (CNNCmut2) behaved similarly, exclusively affecting the G/A variant. This experiment strongly suggests that the binding of SRSF3 is an important determinant of miR-30c expression. Finally, we could recapitulate the observation that SRSF3 binding is limiting for the processing of wild-type pri-mir-30c-1 in an *in vitro* system, supplemented with purified SRSF3 protein (Fig. 6). Firstly, we found that the FLAG-Drosha/FLAG-DGCR8 complexes used for the *in vitro* processing assays contained residual levels of SRSF3 protein **(Supplementary Fig. 5)**. Thus, the relative higher processing of the G/A variant can be explained by the preferential binding of SRSF3 present in the reaction to the G/A variant. Importantly, addition of purified SRSF3 protein increases the Microprocessor-mediated production of WT pre-mir-30c-1 (**Fig. 6a**), whereas addition of purified SRSF3 to the G/A variant (**Fig. 6c**) or to ΔCNNC variants that lack SRSF3 binding sites did not affect the processing activity (**Fig. 6b**, **d**). This is reminiscent of what was observed in MCF7 cells under variable levels of SRSF3.

**Figure 6.**
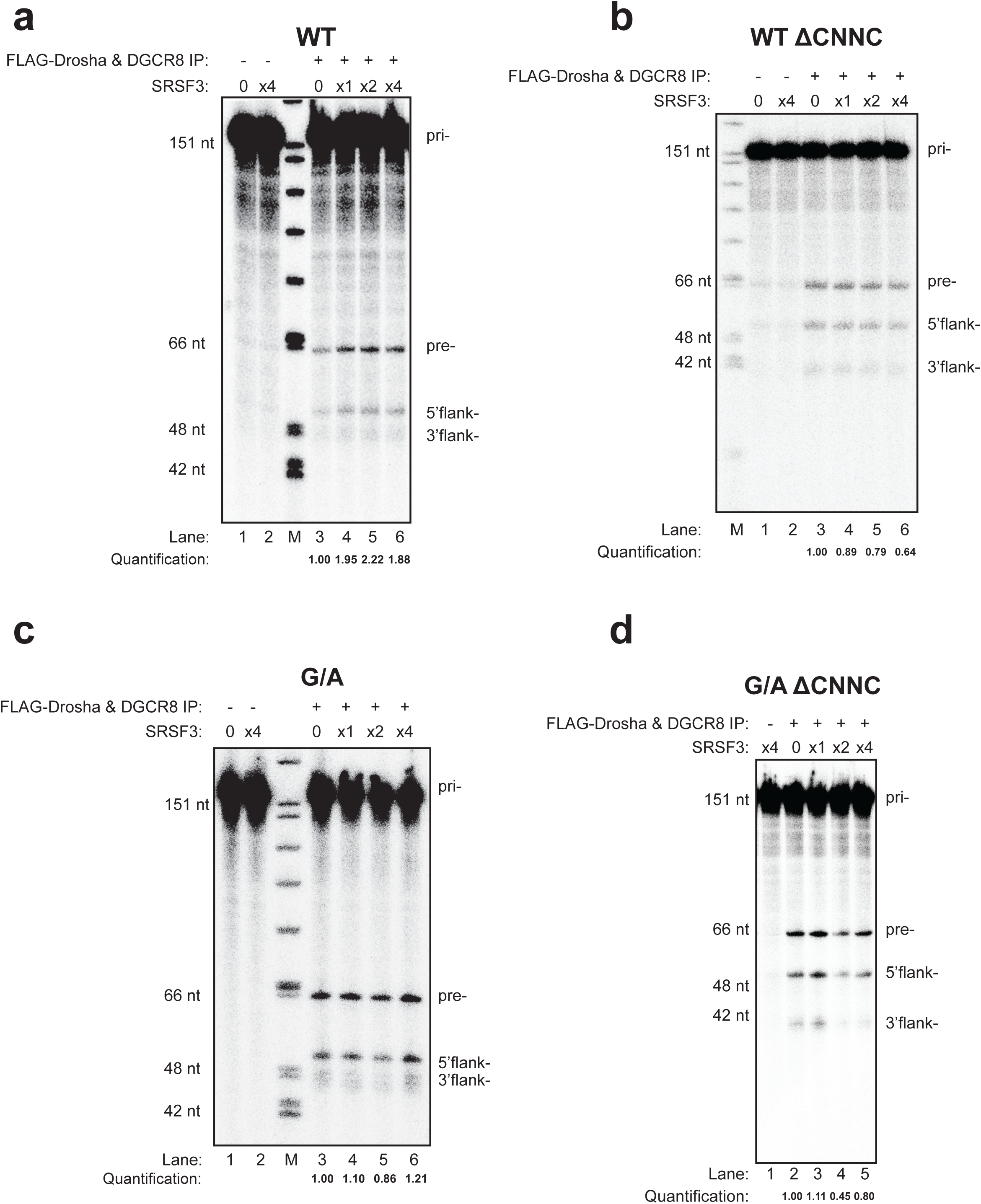
Representative *in vitro* processing assays of radiolabeled shorter pri-mir-30c-1 variants in the presence of immunopurified FLAG-Drosha/FLAG-DGCR8 complexes (+) or FLAG immunoprecipitates (−), with addition of purified SRSF3 protein (1x corresponds to 65nM of protein). (a, b) Processing of WT pri-mir-30c-1 comprising or lacking SRSF3 binding sites (WT and WT ΔCNNC, respectively). (c,d) Processing of pri-mir-30c-1 G/A variant comprising or lacking SRSF3 binding sites (G/A and G/A ΔCNNC, respectively). Quantification of pre-miRNA band intensities are shown below and expressed as the relative intensity normalized to *in vitro* processing assay in the absence of purified SRSF3 protein (lane 3 in each case).

## DISCUSSION

The central role of miRNAs in the regulation of gene expression requires that their expression is tightly controlled. Indeed, the biogenesis of cancer-related miRNAs, including those with a role as oncogenes (‘oncomiRs), or those with tumor suppressor functions is often dysregulated in cancer^13,15,38^. Interestingly, some miRNAs have been shown to display both tumor suppressor and also oncogenic roles, depending on the cell type and the mRNA targets^39^, as was described for miR-221, which exerts oncogenic properties in the liver^40^ but also acts as a tumor suppressor in erythroblastic leukemias^41^. Futhermore, miR-375 has been shown to display a dual role in prostate cancer progression, highlighting the importance of the cellular context on miRNA function^42^.

Despite a more comprehensive knowledge on the role of RNA-BPs in the posttranscriptional regulation of miRNA production, there is only circumstantial evidence on how RNA sequence variation and RNA structure impact on miRNA processing. There are several reports showing that a single nucleotide substitution in the sequence of precursor miRNAs could have a profound effect in their biogenesis. Nonetheless, there is limited information about SNPs in the terminal loop region (TL) of pri-miRNAs. A bioinformatic approach led to the identification of 32 such SNPs in 21 miRNA loop regions of human miRNAs^43^. Some studies have found a correlation between the presence of polymorphisms in pri-miRNAs and expression levels of their corresponding mature miRNAs, affecting cancer susceptibility, as shown for miR-15/16 in chronic lymphocytic leukemia (CLL)^44,45^, miR-146 in papillary thyroid carcinoma^46^ and miR196a2 in lung cancer^47^. Another example is the finding of a rare SNP in pre-miR-34a, which is associated with increased levels of mature miR-34a. This could be of biological significance since precise control of miR-34a expression is needed to maintain correct beta-cell function, thus this could affect type 2 diabetes susceptibility^48^. The emerging picture is that human genetic variation could indeed not only have a role in miRNA function by affecting miRNA seed sequences and/or miRNA binding sites in the 3’UTRs of target genes, but it can also contribute significantly to modulation of miRNA biogenesis^21^.

In this study, we focused on a rare genetic variation found in the conserved terminal loop of pri-mir-30c-1 (G_27_ to A variant) that was found in breast cancer and gastric cancer patients and leading to increased expression of miR-30c^22,23^. There is circumstantial evidence that miR-30c is involved in many malignancies acting as tumour suppressor^49,50^ or as an oncogene^51–53^. In order to understand the mechanism underlying miR-30c deregulation in breast cancer, we investigated how this mutation affects miRNA biogenesis. We show that the G-A substitution in pri-mir-30c-1 directly affects Drosha-mediated processing both *in vitro* as well as in cultured cells (**Fig. 1**, 5, 6 and **Supplementary Fig. 1)**. The conservation of pri-mir-30c-1 sequences across vertebrate species highlights the importance of the primary sequence in the TL, 5´ and 3´ regions (**Fig. 2**), suggesting a crucial role in miRNA biogenesis. Indeed, conserved sequences in TL have been shown to be important for recognition by auxiliary factors^24^ as well as for DGCR8 binding^54^, allowing efficient and accurate miRNA processing. It has also been shown that pri-miRNA tertiary structure is a major player in the regulation of miRNA biogenesis, as observed for the well characterized miR17-92 cluster^55–57^. Here, using SHAPE structural analysis, in conjunction with solvent accessibility analysis by hydroxyl cleavage, we found that the G/A sequence variation leads to a structural rearrangement of the apical region of the pri-miRNA affecting the conserved residues placed at the basal part of the stem (**Fig. 3** and **Supplementary Fig. 2)**. This demonstrates that pri-mir-30c-1 is organized as a complex and flexible structure, with the TL and the basal region of the stem potentially involved in a tertiary interaction. Further work is required to determine the existence of direct contacts between these regions.

Interestingly, we also observed that this RNA structure reorganization promotes the interaction with SRSF3, an SR protein family member that was demonstrated to facilitate pri-miRNA recognition and processing^34^, by recognizing the CNNC motif located 17nts away from Drosha cleavage site. Pri-mir-30c-1 has two overlapping CNNC motifs (residues from +16 to +21) (**Fig. 5b**). Notably, accessibility around this region increased in G/A variant (**Fig. 3**). Furthermore, toeprint and SHAPE assays in the presence of purified SRSF3 protein clearly demonstrated that SRSF3 is specifically recognizing the CNNC motif in a dose-dependent manner (**Fig. 4**).

Altogether, data presented here suggest that binding of SRSF3 to the wild-type sequence is limiting and that the structural reorganization induced by the G/A substitution makes the SRSF3 binding sites more accessible. Taking everything into account we propose a model whereby a genetic variant in a conserved region within the TL of pri-mir-30c-1 causes a reorganization of the RNA secondary structure promoting the interaction with SRSF3, which in turn enhances the Microprocessor-mediated processing of pri-mir-30c-1 leading to increased levels of miR-30c (**Fig. 7**). We conclude that primary sequence determinants and RNA structure are key regulators of miRNA biogenesis.

**Figure 7.**
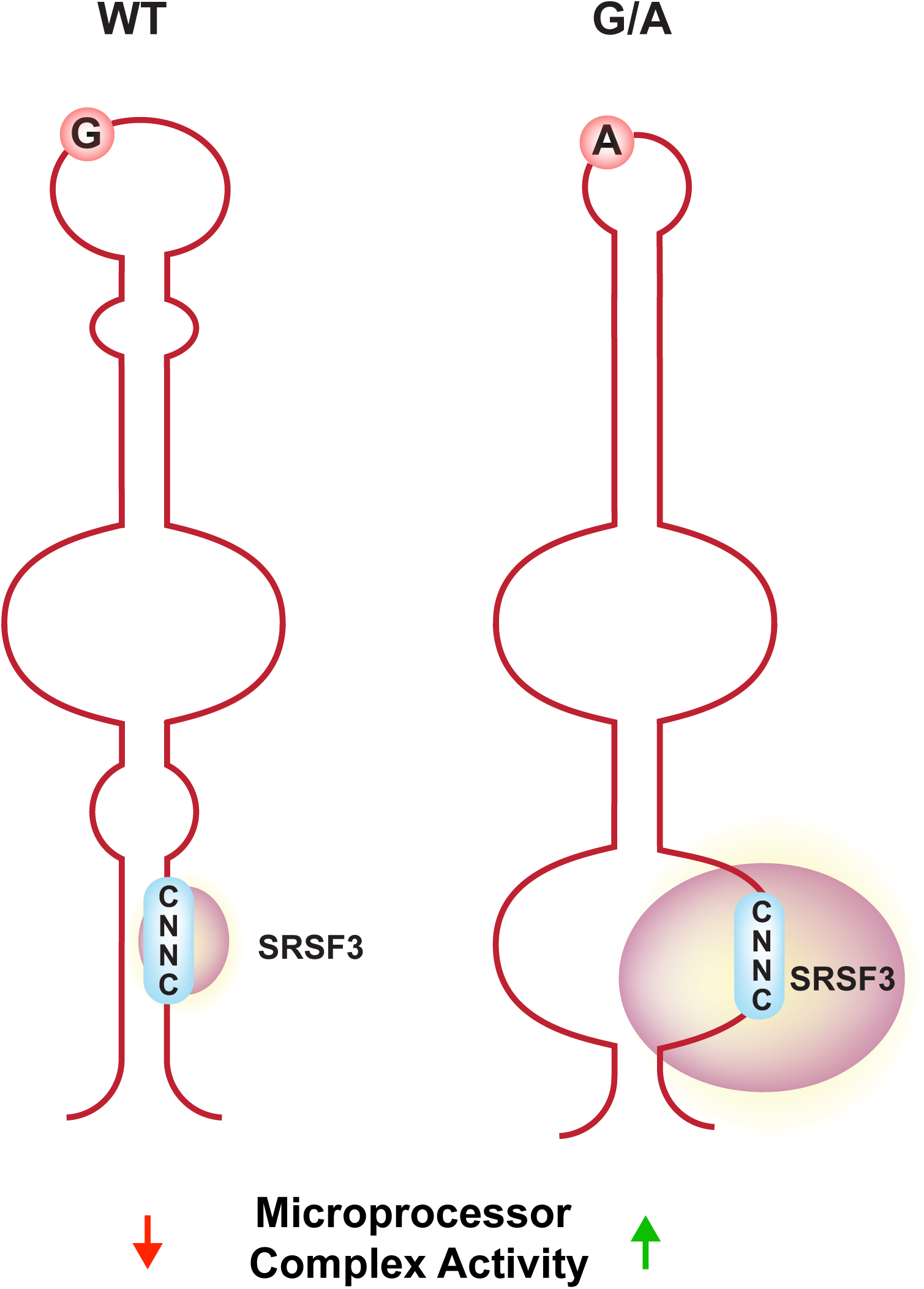
Cartoon depicting a model whereby the genetic variant identified in pri-mir-30c-1 leads to a secondary RNA structure rearrangement that facilitates binding of SRSF3 resulting in increased Microprocessor-mediated processing of pri-miR-30c-1.

## METHODS

Methods and any associated references are available in the online version of the paper

## ACKNOWLEDGEMENTS

We are grateful to David Fitzpatrick and Magdalena Maslon (MRC HGU) for discussions and to Javier Martinez (IMBA, Vienna) and Encarnación Martínez-Salas (CBM, Madrid) for critical reading of the manuscript. This work was supported by Core funding from the Medical Research Council and by the Wellcome Trust (Grant 095518/Z/11/Z).

## AUTHOR CONTRIBUTIONS

N.F., R.A.C, S.M. and J.F.C. conceived, designed, interpreted the experiments and wrote the manuscript. N.F. performed most of the experiments and data analysis. R.A.C. carried out *in vitro* pri-miRNA processing and N.H. performed biochemical experiments. S.M. carried out some of the initial experiments. R.S.Y. provided bioinformatics analysis. J.F.C. supervised the whole project.

## SUPPLEMENTAL INFORMATION

Supplemental information includes 5 figures, Supplemental Experimental Procedures, and Supplemental References and can be found with this article online

**Supplementary Figure 1** Representative *in vitro* processing of shorter pri-mir-30c-1 substrates (153nt) comprising either a wild-type sequence (WT) or bearing the G/A mutation in the presence of immunopurified FLAG-Drosha/FLAG-DGCR8 complexes (+) or FLAG imunoprecipitates (−). Quantification of pre-miRNA band intensities are shown below and expressed as the relative intensity normalized to pre-mir-30c-1 WT variant (lane 3). Intervening lanes were removed from the gel image, which is indicated by the black vertical line.

**Supplementary Figure 2 (a)** Histograms of cleavage intensity versus nucleotide position determined using *in situ*-generated hydroxyl radicals values corresponding to the mean cleavage (+/− SEM) (n=6). **(b)** Predicted secondary structures of WT and G/A variant sequences, as determined by hydroxyl radical reactivity values.

**Supplementary Figure 3** Identification of proteins interacting with WT and G/A pri-miR30c-1 sequences. **(a)** Table shows proteins identified in two independent experiments. **(b)** Validation of the interaction of SRSF3 with the G/A variant sequence by RNA immunoprecipitation followed by Western Blot analysis with specific antibodies.

**Supplementary Figure 4** SRSF3 levels following knock-down or overexpression in MCF7 cells. **(a)** Analysis of SRSF3 mRNA levels. Error bars indicate SEM (n=5). **(b)** Representative Western blot of SRSF3 protein levels.

**Supplementary Figure 5** Detection of SRSF3 protein in FLAG-Drosha/FLAG-DGCR8 complexes used in the *in vitro* processing assays shown on Fig 6 and Supplementary Figure 1. FLAG immunoprecipitates of control cells or cells transfected with FLAG-Drosha + FLAG-DGCR8 were analyzed in a Western blot assay with an anti-SRSF3 specific antibody.

## ONLINE METHODS

### Plasmids constructions

Pri-mir-30c-1 construct was amplified from human genomic DNA by PCR with specific primers 30c1s (5’-CAAGTGGTTCTGTGTTTTTATTG-3’) and 30c1a (5’-GTACTTAGCCACAGAAGCGCA-3’) The PCR product was digested with *EcoRl* and was subsequently cloned into the pCDNA3.1 vector (ThermoFisher). The G_27_ to A mutation was generated by a two-step PCR strategy. First, pri-miR-30c-1 was amplified with a 30Cmut1 oligo (5’-CCTTGAGCTTACAGCTGAGAG-3’) and 30c1s and also with 30Cmut2 oligo (5′-CTCTCAGCTGTAAGCTCAAGG-3′) and 30c1a. Both PCR products, were purified (Qiagen), pooled and used as a template for amplification with 30c1s and 30c1a primers. The resulting PCR product was cloned in pGEMt (Promega). pGEMt G/A plasmid was digested with EcoRI to clone in pCDNA3.1. The CNNC motifs were subjected to site-specific mutagenesis by PCR amplification, as previously described^58^. Briefly, 10 ng of pri-miR30c-1 (WT or G/A) was PCR amplified with the desired mutagenic oligonucleotide (mut1: 5´-CTTCATTTGATGTTTTCCATGGC-3´, mut2: 5′-CTTCTTTTTTTTTTTCCATGGC-3′ or CNNC 5′-CTTCAGATGTTTTCCATGGC-3′)and CNNCs primer (5´-CTGCTTACTGGCTTATCG-3´). The PCR product was cleaned (Qiagen) and used as the 5´-flanking primer in a second PCR with an equal molar amount of primer CNNCa (5´-GATATCTGCAGAATTCACTAG-3´). The product of the second PCR was digested with *EcoR*I (New England Biolabs), purified by agarose gel electrophoresis and ligated to the large *EcoR*I fragments of pri-miR-30c-1 to produce the desired constructs (CNNCmut1, mut2 and ΔCNNC). All the sequences were confirmed by automatic sequencing.

### Cell Culture

MCF7 and HEK 293T cells were grown in high glucose Dulbecco’s modified Eagle’s medium (Invitrogen) supplemented with 10% (v/v) fetal calf serum (Invitrogen) and penicillin-streptomycin (Invitrogen) and incubated at 37°C in the presence of 5% CO_2_. Cells were tested for mycoplasma contamination.

### Transfections

MCF7 cells grown in 24-well plates were transfected with either pri-mir-30c-1 (WT or G/A) constructs, *in vitro* transcribed RNA (0.33μg/10^5^ cells), or oligonucleotides encoding pre-mir-30c (Sigma Aldrich)(5´UGUAAACAUCCUACACUCUCAGCUGUGAGCUCAAGGUGGCUGGGAGAGGGUUGUUUACUCC-3´) using Attractene (Qiagen), following manufacturer’s instruction. pCDNA-3, pri-miR-30a or oligo30a (5´UGUAAACAUCCUCGACUGGAAGCUGUGAAGCCACAAAUGGGCUUUCAGUCGGAUGUUUGCAGC-3´) were used as negative controls in DNA or RNA transfections, respectively. Cells extracts were prepared at 8h or 48h (after RNA/DNA addition) by direct lysis using 100 μl of lysis buffer (50mM Tris-HCl at pH 7.8, 120 mM NaCl, 0.5% NP40). For SRSF3 gene silencing/overexpression MCF7 cells grown in 15 cm dishes were transfected with ON-TARGETplus siRNA (Dharmacon) or pCG. As negative controls, cells were transfected with ON-Targetplus Non-targeting siRNAs and pCG-T7 plasmid, respectively. Cells were split in 24-well dishes 24h after transfection and 24h later transfected with different versions of pri-miRNA constructs. HEK 293Ts were grown to 70% confluency in 6-well plates and then transiently co-transfected with 3μg of FLAG-Drosha and 1μg of FLAG-DGCR8, or 4μg FLAG-empty vector per well. Cells were expanded for 24h, then split to 10cm plates and expanded for a further 24hr before cells were scraped, collected and snap frozen until required.

### SRSF3 purification

Purification of SRSF3 from MCF7 cells was performed as previously described^59^, with minor modifications. Briefly, MCF7 cells were grown to confluence in 15 cm dishes and transient transfected with 15 μg of pCG-T7-SRSF3^60^. Forty eight hours after transfection cell pellets were resuspended in 20ml of ice-cold lysis buffer (50 mM NaP buffer, pH8, 0.5 M Na Cl, 5 mM b-glycerophosphate, 5 mM KF,0.1 % Tween 20, and 1x protease inhibitor cocktail) and sonicated 5 times 30s followed by centrifugation at maximum speed for 20 min at 4°C. After centrifugation the supernatant was loaded into a chromatography column (Biorad) previously prepared with T7 Tag antibody Agarose (Novagen). The flow-through was collected and loaded a second time. The column was washed two times with 10 ml of lysis buffer. Then, eluates were eluted with 10 serial 0.8 ml volumes of elution buffer (0.1 M citric acid, pH 2.2, 5 μm β-glycerophosphate cand 5mM KF) and collected in microcentrifuges tubes containing 200 ul of 1M Tris pH 8.8 mixed and stored at 4°C. The fractions were analysed on SDS page (Invitrogen) followed by Coomassie blue staining. The elutes containing the protein were dialyzed overnight against BC100 buffer (20 mM tris, pH8, 100 Mm KCl, 0.2 mM EDTA pH8 and 20% glycerol), and stored in aliquots at −80°C.

### Western blot analysis

Equal amounts of total protein (determined by Bradford assay) were loaded in 12% NUPAGE gels (Invitrogen) and transferred to cellulose membranes using IBLOT system (Invitrogen). Identification of SRSF3 was performed with a rabbit polyclonal antibody (RN080PW, MBL, Dilution 1:500), followed by a secondary horseradish peroxidase-conjugated antibody and ECL detection (Pierce). Other primary antibodies used in this study were: mouse monoclonal anti-PARP-1 antibody (E-8): sc-74469, Santa Cruz Biotechnology, Dilution 1:500); Rabbit polyclonal anti-DDX17 antibody ((S-17): sc-86409, Santa Cruz Biotechnology, Dilution 1:500); Rabbit polyclonal anti-hnRNP A1 antibody (PA5-19431, Invitrogen antibodies, Dilution 1:500); Rabbit polyclonal anti-TLS/FUS antibody (ab23439, Abcam, Dilution 1:1000).

### *In vitro* transcription of pri-miRNA substrates

Prior to RNA synthesis, pri-mir-30c-1 plasmids (wild-type and G/A variant) were linearized using *Apa*I (New England Biolabs). In addition, shorter pri-mir-30c-1 probes were PCR amplified from pri-mir-30c-1 plasmids (wild-type, G/A, and their respective ΔCNNC) for *in vitro* processing assays (shown on **Fig. 6** and **Supplementary Fig. 1)** using a forward primer harboring a <u>T7-promoter</u> sequence fused to a 19nt sequence complementary to pri-mir-30c-1 (5’-<u>AATACGACTCACTATAG</u>GGCTGATCAACCCTGGACC-3’) and a reverse primer (5’-AGTGGAGACTGTTCCTTCTTC-3’), which when *in vitro* transcribed generated a 153nt (CNNC) or 146nt (ΔCNNC) and then subsequently PCR purified (Qiagen) for *in vitro* transcription. 1μg of DNA or 400ng of PCR product were *in vitro* transcribed for 1-2h at 37°C using 50 U of T7 RNA polymerase (Roche) in the presence of 0.5 mM rNTPs and 20 U of RNAsin (Promega). When needed, RNA transcripts were labeled using (α-^32^P)-UTP (800 Ci/mmol, Perkin Elmer) and following DNAse treatment, unincorporated ^32^P-UTP was eliminated by exclusion chromatography in TE equilibrated columns (GE Healthcare) or PAGE gel purification, followed by phenol/chloroform and ethanol precipitation.

### *In vitro* processing assays

The *in vitro* processing reactions were performed as previously described^61^ with minor modifications. Radiolabeled *in vitro* transcribed pri-mir-30c-1 (50,000 c.p.m.) was incubated with 650μg of MCF7 total cell extract (Fig. 1d), or incubated with FLAG-tagged complexes immunopurified from HEK293T cells transiently coexpressing FLAG-Drosha & FLAG-DGCR8 or FLAG-empty vector control, in the absence or RNase^62^ (**Fig. 6** and **Supplementary Fig. 1)**. Additionally, *in vitro* processing reactions were supplemented with increasing concentrations of immunopurified T7-SRSF3 (**Fig. 6**). The *in vitro* processing reactions were performed in the presence of buffer A (0.5 mM ATP, 20 mM creatine phosphate and 6.4 mM MgCl_2_). Reactions were incubated for 1h at 37°C and treated with proteinase K. RNA was extracted by phenol/chloroform and ethanol precipitation. Samples were resolved in an 8% 1× TBE polyacrylamide urea gel.

### miRNA qRT-PCR

The relative amounts of miR-30c present in total RNA samples were measured using Exiqon miRCURY LNA^TM^ following manufacturer’s instructions. Total RNA from cytoplasmic lysates was isolated using RNAzol (Invitrogen) and RT was carried out with Universal RT microRNA PCR using 500 ng of RNA. A 1/20 dilution of the RT reaction was used for qPCR with a specific microRNA LNA^TM^ primers set and the LightCycler system with the FastStart DNA Master Green II (Roche). The amount of miR-30c detected in the reaction was normalized by a parallel reaction performed with U6 and SNOR48 primers.

### Quantitative RNA co-immunoprecipitation (qRNAco-IP)

The qRNAco-IP was performed as previously described^63^, with several modifications. Briefly, lysates from pri-miR-30c-1 (WT and G/A) transfected cells were pre-cleared with mouse IgG beads followed by incubation with a polyclonal rabbit anti-SRSF3 antibody (MLB). The complexes were pulled-down using protein G beads (Amershan), then treated with proteinase K (Sigma) and RNA was extracted and purified using Trizol (Invitrogen). qRT-PCR was carried out with superscriptIII (Invitrogen) RT using 500ng of RNA as a template and the TL (5’-GGAGTAAACAACCCTCTCCCAGC-3’) and actin (5’-GGTCTCAAACATGATCTGGG-3’) primers. Then a 1/20 dilution of the RT reaction was used for qPCR with appropriate pairs of specifics primers (TL: 5’-CATCCTACACTCTC-3’ and 5’-GGAGTAAACAACCCTCTCCCAGC-3’, actin: 5’-GGGTCAGAAGGATTCCTATG-3’ and 5’-GGTCTCAAACATGATCTGGG-3’). qRNAco-IP values were calculated according to the formula: qPCR value TL mRNA IP/qPCR value TLmRNA total)/ (qPCRvalue control mRNA IP/qPCR value control mRNA total)

### Evolutionary constraint

Evolutionary constraint was quantified for individual nucleotide positions as the number of rejected substitutions, as calculated by the GERP++ algorithm. This data was extracted from the UCSC genome browser, where they had been calculated over their 36-way mammalian genome alignments.

### Phylogenetic conservation

The alignment of 98 vertebrate sequences of pri-mir-30c-1 were retrieved from USCS genome browser. Each nucleotide frequency was plotted in a heat map. RNA structure models were performed using RNAstructure software (http://rna.urmc.rochester.edu/RNAstructureWeb/). The most energetically stable RNA structure according to the program was used to depict the model of pri-mir-30c-1.

### SHAPE analysis

Pri-mir-30c-1 RNA was treated with NMIA (Invitrogen), as previously described^64^. For primer extension, equal amounts of NMIA treated and untreated RNA (0.5 pmols) were incubated with 2 pmol of 5´-end fluorescently-labeled primer (5´-CTAGATGCATGCTCGAGCG-3´). NED fluorophore was used for both, treated and not treated RNAs while VIC fluorophore was used for the sequencing ladder, as described previously. cDNA products were resolved by capillary electrophoresis^64^.

Pri-miR-30c-1-SRSF3 complexes were assembled in folding buffer (100 mM HEPES pH 8, 6 mM MgCl_2_) using 170 nM RNA in the presence of increasing amounts of purified SRSF3 protein (260 and 500 nM). Then, RNA alone or pre-incubated with SRSF3 was treated with NMIA. RNA was phenol extracted and ethanol precipitated and then subjected to primer extension analysis.

### Hydroxyl radical footprinting

Pri-mir-30c-1 RNA was subjected to Hydroxyl radical footprinting, as previously described^65^. Briefly, 1.7 pmol of RNA was denatured and folded in folding buffer (40 mM MOPS pH 8.0, 80 mMKOAc, and 0, 0.5 or 6 mM MgCl_2_). Samples were incubated with 1 μL of the Fe(II)-EDTA complex, 1 μL of sodium ascorbate and 1 μL of hydrogen peroxide for 30 s at 37 °C. Fe(II)-EDTA (7.5 mM Fe(SO_4_)2(NH_4_)2·6H_2_O and 11.25 mM EDTA, pH 8.0), 0.3% hydrogen peroxide and 150 mM sodium ascorbate solutions were freshly prepared. As a control a lacking Fe(II)-EDTA reaction was performed. Samples were quenched and precipitated by addition of one-third of 75% glycerol, 1 μL of 20 mg·mL^−1^ glycogen, 1 μL of 3 MNaCl, 2 μL of 0.5 M EDTA and 2.5 volumes of ice-cold ethanol. RNAs were re-suspended and reverse-transcribed using fluorescent primers as described for SHAPE reactivity. cDNA products were resolved by capillary electrophoresis.

### SHAPE reactivity and Hydroxyl radical cleavage data analysis

SHAPE electropherograms of each RT were analized using the quSHAPE software^66^. Then, data from 10 independent assays were used to calculate the mean of SHAPE reactivity. To minimize the technical variation, each RT stop quantitative value was normalized to the total intensity of the average of 10 highly reactive nucleotides in the reaction. Grubbs' test was used to identify outliers. To obtain SHAPE differences between pri-mir-30c WT and GA or Free RNA and RNA-SRSF3 complexes, the SHAPE reactivity values obtained in WT or free RNA were subtracted from the reactivity values obtained in GA or in the presence of the protein. The statistical analysis was performed by Mann-Whitney *U* test. Only nucleotides with absolutes differences larger than 0,25 and statistically significant were considered. Hydroxyl radical cleavage intensities of each reaction were also analized using quSHAPE software^66^. Then, data from 6 independent assays were used to calculate the mean of hsa-pri-mR-30c WT or GA cleavage. Grubbs’ test was used to identify outliers. Buried regions were defined as zones with more than two consecutives nucleotides having a reactivity (R) smaller than the mean of all reactivity, whereas exposed regions are those with more than two consecutive nucleotides having R larger than the mean of all reactivity.

### Toeprint assays

This assay carried out following the protocol previously described^35^, with minor modifications. Briefly, pri-mir-30c-1: SRSF3 complexes were assembled as described for SHAPE analysis. After protein incubation samples were subsequently subject to primer extension using fluorescent primer (NED) 5’CTAGATGCATGCTCGAGCG-3.pri-miR-30c. Primer extension products were extracted with phenol, and ethanol precipitated pellets were resuspended in 5 μl of HI-Di ^TM^ formamide (ThermoFisher),which included 0.5 μl of GeneScan™ 500 Liz^TM^ dye size standards. The products were separated by capillary electrophoresis and analyzed by GeneMarker software.

### RNA chromatography

RNase-assisted RNA chromatography with RNAse A/T1 was performed, as described^28^, using *in vitro* transcribed pri-mir-30c-1 and total MCF7 cell extracts. RNA-bound proteins were separated using 4-12% NUPAGE bis-tris system (lnvitrogen) and individual lanes were subjected to mass spectroscopy (BSRC Mass Spectrometry and Proteomics Facility, University of St Andrews). Results were confirmed by Western blot analysis with anti-SRSF3 (MLB), anti-PARP-1 (Santa Cruz), anti-DDX17 (Santa cruz), anti-hnRNPA1 (Thermo Scientific) and anti-FUS (abcam) specific antibodies.

